# Efficient encoding of large antigenic spaces by epitope prioritization with Dolphyn

**DOI:** 10.1101/2023.07.30.551179

**Authors:** Anna-Maria Liebhoff, Thiagarajan Venkataraman, William R Morgenlander, Miso Na, Tomasz Kula, Kathleen Waugh, Charles Morrison, Marian Rewers, Randy Longman, June Round, Stephen Elledge, Ingo Ruczinski, Ben Langmead, H Benjamin Larman

## Abstract

We investigated a relatively underexplored component of the gut-immune axis by profiling the antibody response to gut phages using Phage Immunoprecipitation Sequencing (PhIP-Seq). To enhance this approach, we developed Dolphyn, a novel method that uses machine learning to select peptides from protein sets and compresses the proteome through epitope-stitching. Dolphyn improves the fraction of gut phage library peptides bound by antibodies from 10% to 31% in healthy individuals, while also reducing the number of synthesized peptides by 78%. In our study on gut phages, we discovered that the immune system develops antibodies to bacteria-infecting viruses in the human gut, particularly *E*.*coli*-infecting *Myoviridae*. Cost-effective PhIP-Seq libraries designed with Dolphyn enable the assessment of a wider range of proteins in a single experiment, thus facilitating the study of the gut-immune axis.

## Introduction

The human gut microbiome is a critical determinant of human health. However, the mechanisms underlying the interactions between the host and the diverse microorganisms in the gut, including bacteria, fungi, phages, archaea, and other members of the microbiota, remain largely unknown. Gut phages, which infect bacteria in the gut, are increasingly recognized as important contributors to the host-gut axis. These viruses have even been described as the “puppet masters” of gut bacteria (Camarillo-Guerrero et al. 2021).

The immune system, which defends against foreign invaders and protects tissue homeostasis, plays a major role in gut related health. Secreted host antibodies may, for instance, directly impact the composition of the bacterial population in the gut by neutralizing specific gut phages. Exhaustively characterizing immune response to gut phages requires studying a large number of potential immune targets. Camarillo-Guerrero et al., recently published a database of gut phage sequences, most of which have not been previously described. Though it is smaller than that of gut bacteria, the gut phage proteome is still too vast to be represented via traditional synthetic means.

Phage ImmunoPrecipitation Sequencing (PhIP-Seq) is a technique for profiling the reactivity of an individual’s antibody repertoire to a wide range of antigens. This technique involves designing peptides that tile across proteins, synthesizing oligo libraries that encode the peptides, and cloning the oligo libraries into a phage display vector. Phage display and immunoprecipitation are used to test serum samples for antibody binding to all peptides in parallel, since DNA sequencing is used to determine the relative abundance of the immunoprecipitated population. Protein reference sequences from public databases typically serve as the basis for these phage display libraries that normally consist of 56 to 90 amino acid long peptides. PhIP-Seq libraries have been designed to span the human proteome (Larman et al. 2011), common viruses (Xu et al. 2015), allergens (Monaco et al. 2021), selected gut bacteria (Vogl et al. 2021), and protein toxins (Angkeow et al. 2022), leading to novel insights into health and disease.

To date, PhIP-Seq libraries have been designed using Pepsyn, a software tool that performs uniform peptide tiling across proteins (Mohan et al. 2018). Pepsyn has been typically used to generate peptides that overlap by half the tile size, which results in roughly double coverage of the input proteome. Representing the gut phage proteome in this manner would be intractable which led us to develop a new method for creating more compact PhIP-Seq libraries.

To develop an efficient peptide library, it is necessary to selectively display antibody epitopes, i.e., protein regions that are bound by antibodies. As the study of gut phages remains limited, they are currently underrepresented in databases such as the Immune Epitope Database (IEDB) (Vita et al. 2019). However, it has been recently reported that epitopes recognized by many individuals, known as public epitopes, contain amino acid sequence features that are important for interactions with germline-encoded antibody domains (Shrock et al. 2023). This suggests that we can distinguish peptide sequences more likely to contain epitopes by their amino acid composition, which defines the primary and partially secondary structure of proteins.

Our compact library design method, named Dolphyn, employs two components. The first is a binary machine-learning classifier, trained on public epitope reactivity, for prioritizing small peptides according to their probability of acting as epitopes. The second component is a new strategy for combining multiple regions of a protein into one peptide, allowing for the simultaneous testing of three potential epitopes in a single synthesized oligo. To demonstrate its utility, we used Dolphyn to create a PhIP-Seq peptide library and profiled gut phage proteome antibodies of healthy individuals.

## Results

Traditionally designed peptide libraries tend to contain only a small proportion of reactive peptides. Figure 1A shows the proportion of peptides frequently bound by serum antibodies, with the highest proportion (15.4% per 100 individuals) found in the VirScan library (Xu et al. 2015). The low fraction of reactive peptides even in libraries dominated by human pathogens highlights that such library designs may be significantly improved.

**Figure 1.**
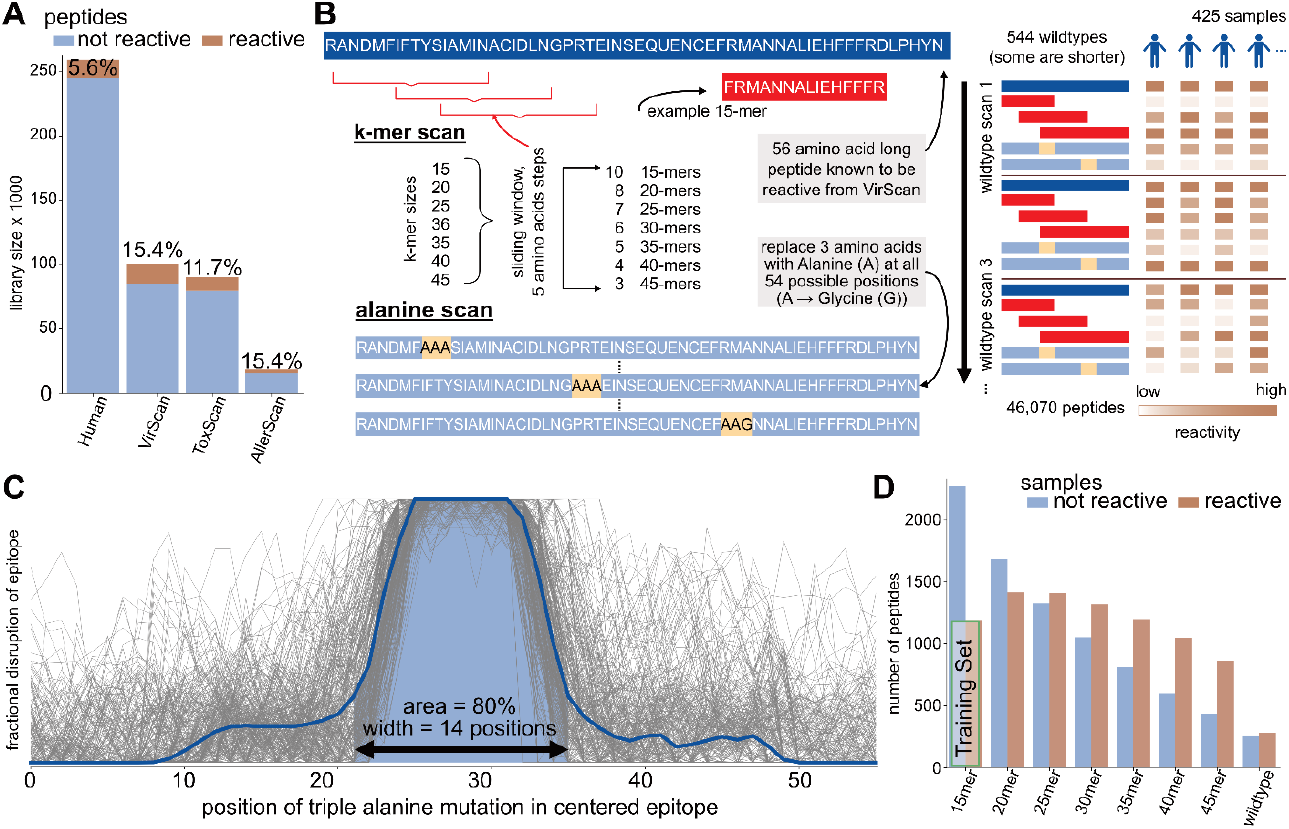
Antibody epitope analysis using programmable phage display of peptide libraries **A** Complexity and reactivity of previously published peptide libraries. Bars show number of peptides included in each library and the percentage of peptides that are reactive in at least 1 of 100 randomly selected samples from the VRC cohort. **B** The Public Epitope Data Set (PEDS) includes a k-mer scan and an alanine scan of 544 virus-derived 56 amino acid long immunodominant peptides in 59 individuals (425 samples). The k-mer scan consists of k = 15 to 45 amino-acid-long sub-peptides of the wildtypes, starting every 5 amino acids. The alanine scan consists of modified versions of the wildtype peptide where triplets of amino acids were replaced with three alanines. Wildtype alanines are replaced by glycines. The library includes one peptide for each modification at each position of a wildtype. **C** Compilation of alanine scans from reactive peptides and individuals. Each grey line is the difference of the alanine peptide at that position to the wildtype reactivity in one individual. Only lines indicating a single epitope were included and shifted to the center. **D** Summarized k-mer scans. A peptide is considered reactive if more than one percent of samples react to it. The “Training Set” indicates those peptides used in the prediction model introduced in Figure 2.

We propose a new method for designing PhIP-Seq libraries for proteomes that are too large to tile exhaustively with Pepsyn. Our method effectively compresses the PhIP-Seq library to a practical size, with minimal loss of sensitivity to detect protein-level reactivity.

### Epitopes contained in reactive peptides

To investigate public epitopes, we selected highly immunogenic peptides (wildtype) for examination in a cohort of 59 individuals (425 samples). The resulting Public Epitope Data Set (PEDS) profiles individual immune responses to two types of sub-peptides of varying lengths, the *alanine scan* and the *k-mer scan* (Methods and Fig 1B).

The alanine scan library of peptides known to contain at least one epitope, contains replicas of the same reactive peptide but replacing each overlapping amino acid 3-mer with three alanines. Alanine substitutions can interrupt antibody reactivity and reveal the location of an epitope within a longer peptide sequence. Figure 1C normalizes, centers, and overlays this information for all individuals eliciting reactivity. 80% of the mean reactivity curve spans 14 amino acid positions, suggesting that most linear public epitopes can be captured by peptides of this length.

Figure 1D presents the results of the k-mer scan to assess reactivity for various peptide lengths. As expected, the number of non-reactive peptides increased with shorter peptide lengths. Wildtype peptides that are reactive may contain one or more epitopes, whereas the shorter sub-peptides derived from the wildtype peptides are likely to contain only one epitope. Shorter peptides are also unlikely to contain many excess amino acids outside of a reactive epitope, as compared to the 56 amino acids long peptides. So, the 15-mers were selected to serve as a dataset for training a binary classifier to determine whether or not a peptide includes an epitope.

### Binary classification of epitope peptides

Approaching epitope prediction as a binary classification problem, we trained a random forest (RF) classifier using peptides that are reactive in many individuals and an equally sized set of peptides that are not reactive in the PEDS cohort. Our model and training data are freely available on GitHub. We provide models for peptides of amino acid lengths 15 to 45.

Fig 2A shows that our classifier fits the data well (AUC 0.99 on the whole dataset, training set was 95%), with the test-set (5%) AUC of 0.69. The higher out-of-bag (OOB) AUC of 0.76 is likely due to existing peptide sequence similarity in the training data, while the test set was selected to avoid sequence similarity to better measure general prediction capability.

**Figure 2.**
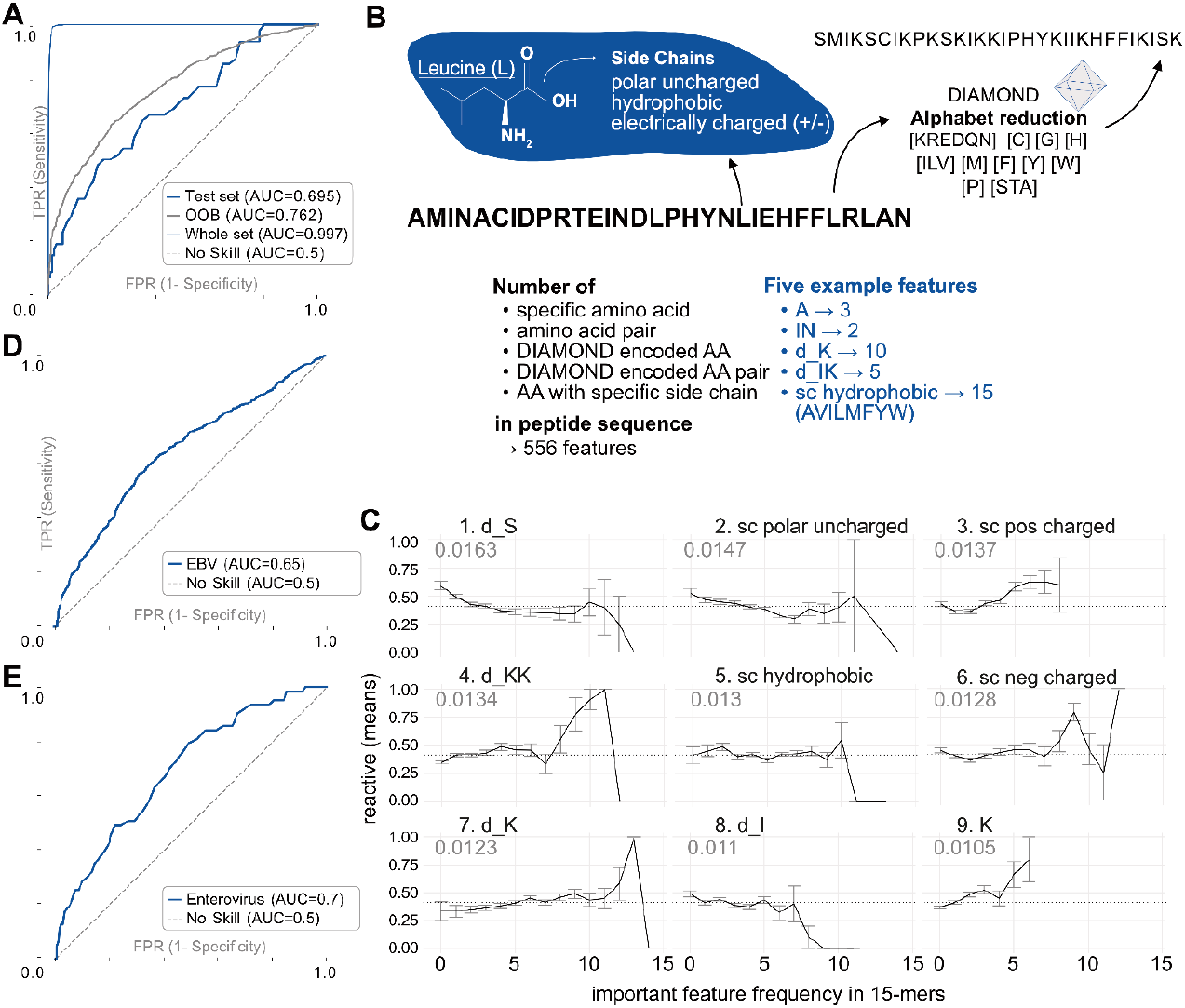
Binary classification of 15-mer peptides containing an epitope. **A** ROC curves for random forest (RF) test and training sets. 5% of the training data was split off as a test set, with no overlap of wildtypes between the sets. **B** Various peptide features were used in a RF model, such as the frequency of natural and DIAMOND encoded AA and AA-pairs, and AA-side chain properties; 556 features in total. **C** Top nine most important features in RF model. The x-axis indicates the frequency of the AA sequence feature (max 15/15) and the y-axis shows the proportion of reactive individuals in the VRC cohort to each peptide in the training set. “d_” indicates a DIAMOND encoded amino acid. The number indicates the mean decrease in impurity, an importance score of a feature. **D** ROC curves on independent 56-mer *Epstein-Barr virus (EBV)* peptides, tested on the VRC cohort. **E** ROC curve of predicted 15-mer peptides in an *Enterovirus* peptide library, tested on the DAISY cohort.

The 556 features used for the model (Fig 2B) included frequencies of amino acids, amino acid pairs, as well as frequencies of classes of amino acids defined by the DIAMOND alphabet reduction (Buchfink, Xie, and Huson 2015) and side-chain type. We found that amino acid frequencies were informative regardless of whether the model included their positions along the peptide.

We discovered the most important features in predicting if a 15-mer contained an epitope to be the amino-acid side-chain frequencies. All four side-chain types were among the nine most important features (Fig 2C). Most crucial appears to be the number of polar uncharged amino acids in the peptide. For instance, Figure 2C illustrates that if a 15-mer peptide contains between 5 to 8 amino acids with a positively charged side chain, it has a higher likelihood of containing an epitope.

The frequency of amino acids in the DIAMOND serine group (including threonine and alanine) is the most important feature. Lysine (K) frequency is also important, as is its DIAMOND group (including arginine, glutamic acid, aspartic acid, glutamine and asparagine), and K-DIAMOND pairs. Interestingly, a recent, independent study also found lysine an important feature of epitopes. A germline encoded feature of antibodies called the “GRAB” motif was described as playing an important role in recognizing public epitopes (Shrock et al. 2023). In humans, these epitopes enrich lysines on their borders if recognized by antibodies using a lambda light chain.

To assess performance on an independent dataset, we collected a set of Epstein-Barr-Virus (EBV) peptides that had been previously screened in a study by Monaco et al. (2021). None of the sub-sequences of this virus had been encountered by our classifier during training. Given the high prevalence of EBV infection worldwide (Arvin et al. 2007), this dataset provided a valuable ground-truth for evaluating the presence of epitopes in these peptides.

Since the EBV peptides were 56 amino acids long, whereas our RF model was trained on 15-mers, we evaluated all possible 15-mers within the 56-mer peptides and used their mean to generate the ROC curve shown in Figure 2D. Despite the necessary transformation to adopt the RF model, the model predicted antibody epitopes with an AUC of 0.65.

We then used the RF model to predict epitopes from 7 Enterovirus strains and selected 757 peptides with low and high probability. For evaluating antibody binding to these peptides with 55 human samples, we defined a peptide as reactive if at least one of these demonstrated reactivity. The ROC curve (AUC = 0.7) is shown in Figure 2E, confirming that the model can distinguish epitopes from non-reactive peptides based on amino acid composition.

### Dolphyn: novel algorithm for peptide library design

Simple tiling methods like Pepsyn (Larman et al. 2011) divide a protein into peptides of equal length with some overlap (Fig 3A). This approach wastes resources on synthesizing, cloning, and sequencing peptides that are not reactive. To improve efficiency, we developed a novel algorithm called Dolphyn that selects and combines peptides with a high probability of eliciting antibody reactivity.

**Figure 3.**
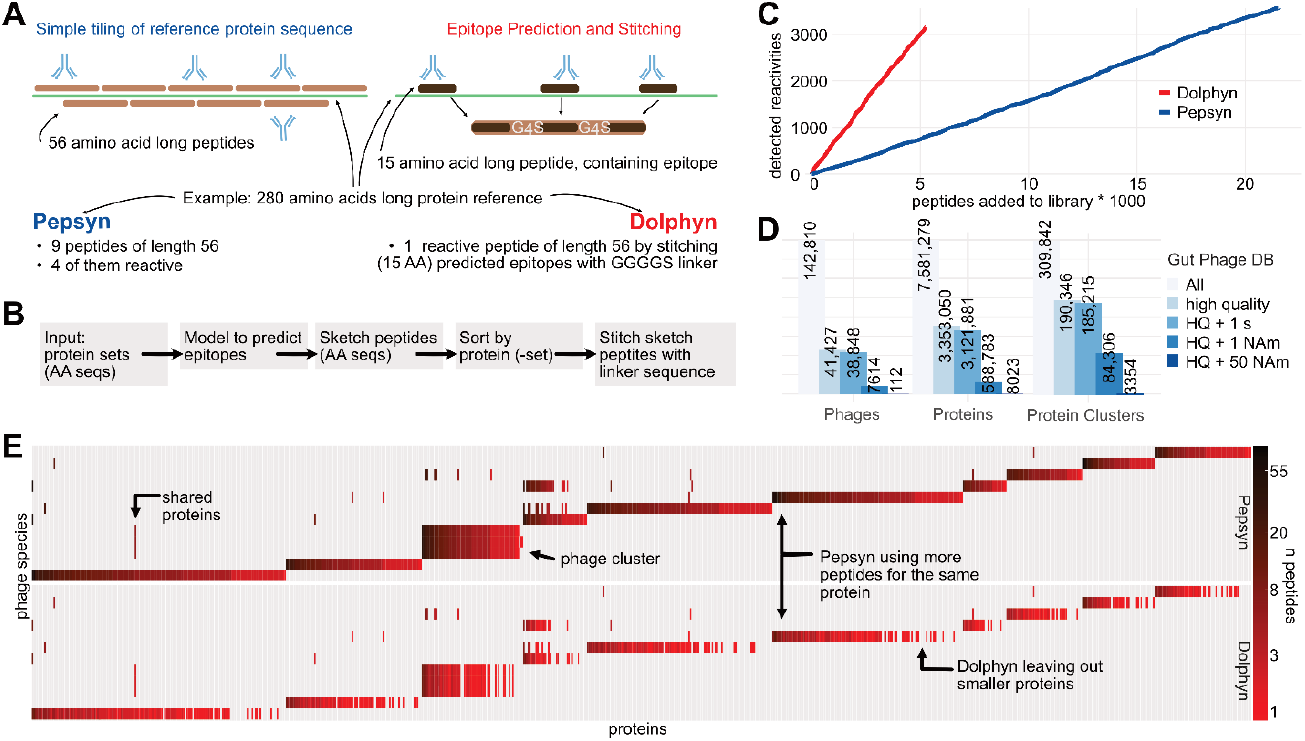
The Dolphyn library design algorithm. **A** Pepsyn tiles the protein sequence uniformly with fixed-size, overlapping peptides. Only a fraction of the peptides is reactive to antibodies. Dolphyn selects a smaller number of short peptides per protein, favoring peptides more likely to be epitopes and stitching them into one composite tile using G4S-linkers to separate the epitopes. **B** Dolphyn Workflow. The package includes modularized steps, e.g. including the RF model (Fig 2). **C** Cost-effectiveness of library design. As more peptides are included, i.e., synthesized and sequenced (x-axis), more immune responses were detected in the test cohort (y-axis). Dolphyn peptides are ordered by their mean prediction value of the contained epitopes, from highest to lowest. Pepsyn peptides are ordered randomly. **D** The metadata analysis of GPD-included phages, proteins and protein clusters reveals sub-sets of genome quality and abundance in North American metagenomic samples. **E** Library composition with Pepsyn versus Dolphyn. 12 randomly selected phage species (y-axis) were represented by Pepsyn or Dolphyn. Proteins are left out when fewer than three epitopes are predicted.

Dolphyn predicts whether each 15-amino-acid sub-peptide (15-mer) of a protein contains an epitope using the RF model described above that was trained on our public epitope dataset (Fig 2). For each protein, Dolphyn selects the three non-overlapping 15-mers with the highest epitope probability (Fig S1). For long proteins, multiple sets of three 15-mers are selected if the probability of containing an epitope is greater than 0.5. These three are combined using Dolphyn’s “stitching” step and separated via a flexible and inert linker sequence GGGGS (see Fig S2 for evaluation of different linkers). The resulting peptide is used to evaluate antibody reactivity at the protein level.

Dolphyn is available as a modular Python package, with parameters controlling epitope peptide length, the linker sequence, the probability cutoff, the training data for the classifier, and the classifier itself (Fig 3B). For example, users can replace the classifier with Immune Epitope Database (IEDB) epitopes or their own classifiers.

To evaluate the efficiency of Dolphyn and Pepsyn libraries based on cost-effectiveness, we accumulated the number of reactivities detected per new peptide added to the library (Fig 3C). In order to detect the same number of immune responses as a Dolphyn library, a Pepsyn library requires about three times the number of peptides.

### Compressing the gut phage database for antibody profiling

(Camarillo-Guerrero et al. 2021) constructed the gut phage database (GPD), which contains 142,810 phage genomes derived from metagenomic stool samples. Roughly one third are marked “High Quality” and 5% were detected in samples from individuals marked as “North American” (Fig 3D). GPD also contains reference amino acid sequences for phage proteins. The authors clustered all proteins at a 95% similarity threshold, resulting in a number of clusters equal to 4% the number of proteins, indicating that the GPD contains many homologous proteins.

We used the GPD reference to design a phage display library for antibody profiling and evaluating Dolphyn. We considered only high-quality phages that appeared in at least one North American individual, leaving 84,000 protein clusters. A Pepsyn-designed library that tiles these clusters would contain over 480,000 56 amino acid long peptides, whereas a Dolphyn designed library requires only about 100,000 peptides (Table 1).

**Table 1.** Peptide library design statistics for Gut Phage Database protein sets.

|  | Pilot Library |  | North American Phageome |  |
| --- | --- | --- | --- | --- |
| **Phages** | 112 |  | 7614 |  |
| **Proteins** | 8023 |  | 588,783 |  |
| **Protein Clusters** | 3354 |  | 84,306 |  |
| **AA length of representative proteins concatenated** | 750,776 |  | 15,637,136 |  |
| **Pepsyn tiles + Coverage** | 23,745 | 1.77 | 484,761 | 1.74 |
| **Dolphyn tiles + Coverage** | 5,266 | 0.32 | 106,762 | 0.31 |
| **Compression ratio (Dolphyn/Pepsyn)** | 0.22 |  | 0.22 |  |

To compare Pepsyn versus Dolphyn library performance, we created a pilot library by selecting the 112 phages most prevalent in North American individuals. We selected one representative from each protein cluster present in these phages. Using Pepsyn’s standard tiling strategy, these proteins are covered 1.77 times using 23,745 56-mers (with 28 amino acid overlaps), whereas Dolphyn covers a third of the proteome, using 5,266 56-mers (Table 1). Dolphyn therefore compresses these phage proteomes by nearly five-fold over the traditional approach. In the design of the full gut phage database proteome library, we observe a similar compression.

Figure 3E presents the protein composition of 12 selected phages from the pilot library. Heatmap colors represent the number of peptides in the library for each protein. Many proteins are shared across phages, and phages within GPD-defined phage clusters share most proteins. Dolphyn omits some of the smaller proteins where the number of potential epitopes required for efficient sketching is not sufficient. Consequently, these proteins are not represented in a Dolphyn-designed library, which is one limitation of the approach.

### Effect of stitching on antibody detection sensitivity

Dolphyn creates stitched peptides by combining potential epitopes from the same protein. Specifically, we used the 15-mer with the highest probability at the first position, followed by the second and third highest probabilities at positions two and three, respectively. In larger proteins that required compression onto multiple peptides, Dolphyn distributes the highest probability epitopes over the first positions of each stitched peptide so as to maximize the total number of independently reactive peptides per protein.

Our pilot library includes both individual 15-mers and their corresponding stitched versions. Figure 4A shows reactivity data from these peptide sets for four representative samples, where two or more peptides were reactive. We observe that only one individual 15-mer was typically reactive, with a preference for the higher probability epitopes (Fig 4B). The log hit-foldchange values for the stitched version were similar to those of the reactive individual 15-mer (Fig 4A), indicating effective representation of the epitope.

**Figure 4.**
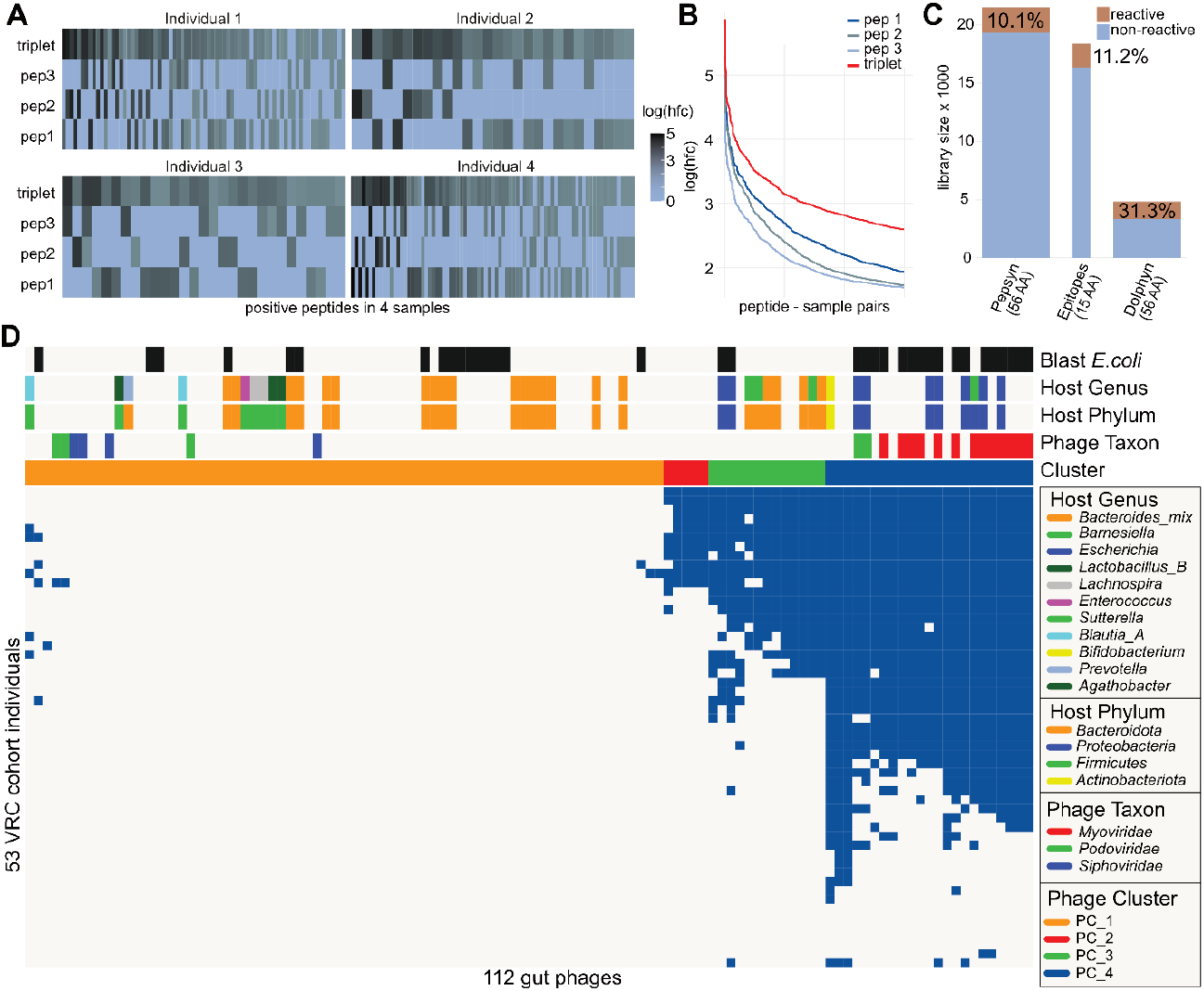
Healthy individuals’ antibody reactivity to 112 gut phages in pilot library. **A** Relationship between reactivities of stitched Dolphyn peptides and unstitched predicted epitope peptides. For four samples, all peptide quartets (three 15-mers and their stitched version) are shown where two or more peptides show reactivity. Peptide sets are ordered by highest to lowest reactivity from left to right. **B** The 15-mer peptides were stitched in the order of their probability score on the combined peptide, starting with the highest at position one. Accordingly, a difference in mean log reactivity score can be observed. Peptides are ordered by reactivity score per sample on the x-axis. **C** The phage proteomes in the pilot library were represented using three different approaches, Pepsyn, predicted epitopes and Dolphyn. The Pepsyn library with regularly spaced 56-mer probes and is the largest but least reactive. The predicted epitopes alone achieve a similar proportion of reactive peptides despite the probes being small (15-mers, depicted by width of bar). The Dolphyn library achieves a greater proportion of reactive probes than the alternatives. **D** Anti-phage antibody reactivities in healthy individuals, depicted as binarized PhARscores, a phage-level aggregate reactivity score. The annotation on the top shows predicted phage properties and phage clusters.

### Reactivity of Dolphyn libraries

We profiled plasma samples from 51 healthy individuals using the three pilot sub-libraries. The Dolphyn library contained a three times higher ratio of reactive peptides (log(hfc)>0 in at least one sample) compared to the Pepsyn library, in which 90% of the peptides were found to be non-reactive (Fig 4C). Individual predicted epitopes displayed only a slightly higher ratio of reactive peptides compared to Pepsyn. However, it should be noted that these peptides are only 15 amino acids long versus the 56 amino acid long Pepsyn peptides and are therefore expected to harbor fewer public epitopes on average.

### Immune response of healthy individuals to gut phages

We then explored the immune response to gut phages in the 51 healthy individuals (Fig 4D). Using a phage-level aggregate metric (PhARscore, Methods), we detected antibody reactivity to a cluster (PC_4) of highly reactive phages in most individuals. Phages in this cluster that had a GPD-predicted taxon belonged to the *Myoviridae* phylum. The predicted phage hosts in this cluster are primarily *Proteobacteria*, especially *E*.*coli*.

As the GPD does not provide predictions of phage taxonomy or host for all phages, we used BLAST to add annotations to the phage genomes. The “Blast *E*.*coli*” heatmap annotation indicates phages with genomes that had an alignment to an *E*.*coli* reference in the NCBI nt database. We assume that these alignments largely correspond to prophage sequences that have integrated into their host bacterium.

### Dolphyn libraries recover observations made with Pepsyn

We confirmed that *E*.*coli* phages and *Myoviridae*-annotated phages elicited a stronger immune response compared to other phages via Wilcoxon test. The mean PhARscore (Methods) for each phage across all samples is significantly higher in all three annotations for both library designs. (Fig 5A)

**Figure 5.**
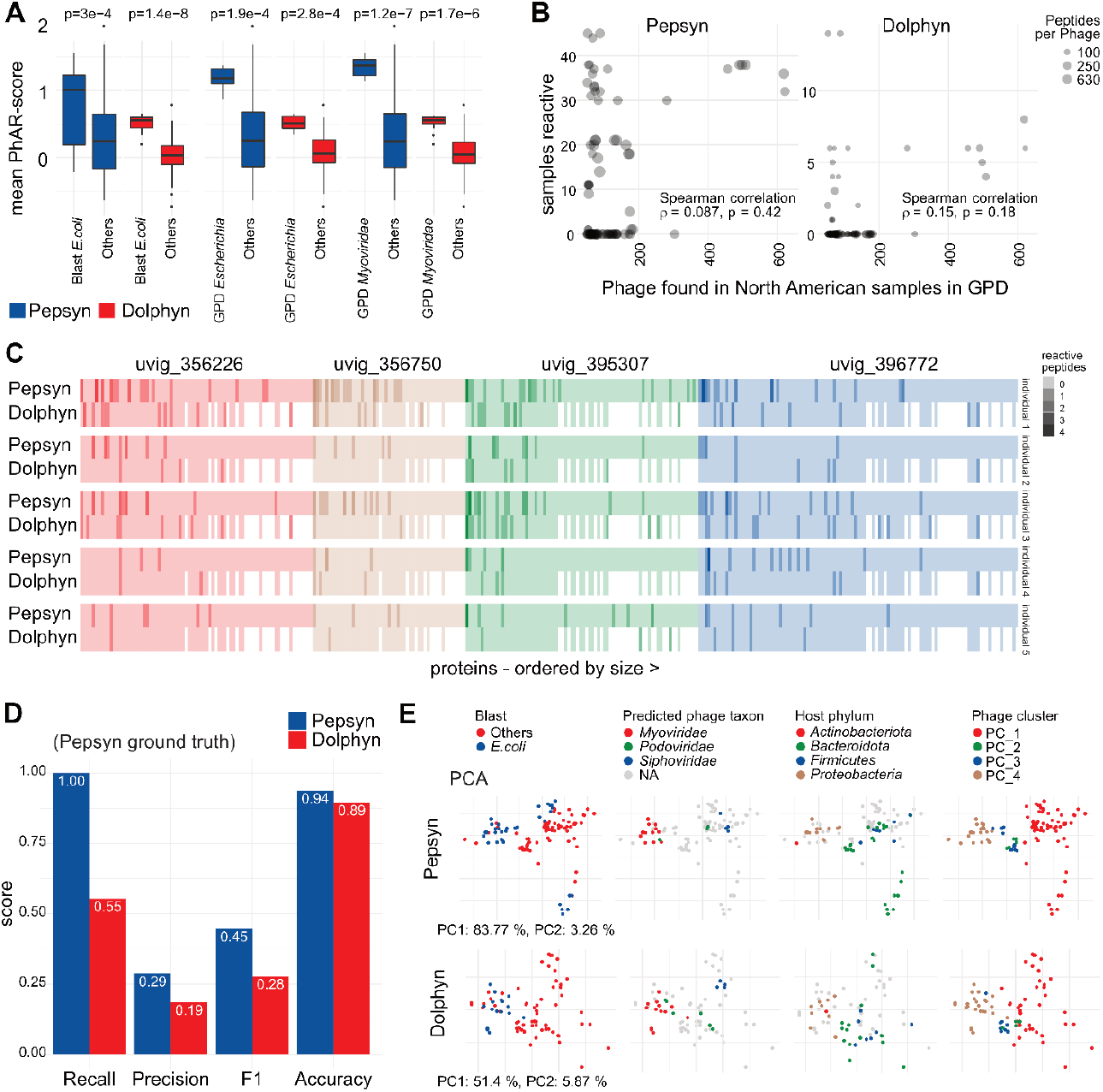
Comparing Dolphyn and Pepsyn results **A** A significant difference in *E. coli* phage reactivity can be observed in mean PhARscore of Pepsyn and Dolphyn designed libraries (Wilcoxon test). **B** The phages detected in more samples in the GPD (x-axis) were also reactive in more individuals in our cohort (y-axis). Phages were only considered if present in both library designs. The spearman correlation is calculated based on reactivity. **C** Phage proteins from phages with PhARscores indicating high reactivity are displayed in sequential order (no shared proteins) for 5 individuals. Note that the Dolphyn library contains less peptides per protein and has consequently a lower number of reactive peptides per protein. **D** Protein discovery power. Using reactivities to Pepsyn proteins as ground truth, performance metrics are shown for both Pepsyn and Dolphyn peptides in recovering protein by sample reactivities. **E** Principal component analysis (PCA) clustering of phages colored by different annotations. The PCs of phages according to the sample-reactivity vectors per phage are similar. Phages with same annotations and clustering in the heatmap (Fig 4) group together independently of the library design.

The GPD is derived from a large set of metagenomic samples, which enabled Camarillo-Guerrero et al. to identify phage presence in those samples. We calculated the prevalence of phages in North American individuals and created the pilot library that includes all phages detected in 50 or more North Americans. The few highly prevalent phages appear to elicit an immune response more commonly across all samples in our cohort, which includes only North American individuals. (Fig 5B)

We investigated what protein level targets drive the immune response to the highly reactive phage cluster in healthy individuals. Figure 5C displays the number of reactive peptides in the proteins of the four most reactive phages in five individuals with the most robust antibody responses according to their PhARscores. We observe that some proteins are detected by several individuals’ antibodies, but overall, the individuals exhibited distinct immune response profiles. The proteins are ordered by size, and a higher number of reactive peptides is expected and observed for larger proteins (towards the left).

Dolphyn-designed libraries demonstrate similar discovery power (accuracy) for identifying protein antibody targets, as shown in Fig 5D. Dolphyn peptides only recall a bit more than half the proteins that are reactive in Pepsyn partially because Dolphyn does not include some proteins. However, Dolphyn was able to identify some proteins that were not reactive in the Pepsyn library and considering this as a “fair” ground truth, Dolphyn achieves higher performance (Fig S3).

The pilot phage library designed with Pepsyn produced several key observations that were recapitulated by the Dolphyn library. The Principal Component Analysis (PCA) shown in Figure 5E is based on sample PhARscores. Color is used in the plot to highlight phage attributes. The rightmost plots show the four identified phage clusters of the heatmap, initially found using the Pepsyn-designed library (Fig 4D). We observed that the Dolphyn-designed library preserved the same clustering for our cohort.

## Discussion

Here we introduce the Dolphyn algorithm for efficiently converting large antigenic spaces into tractable peptide libraries for antibody profiling applications. The algorithm reduces the number of peptides in a library for a given proteome by 78% and triples the reactivity per peptide, as compared to uniform protein tiling.

Dolphyn employs a random forest model which aids in selecting peptides likely to contain an epitope based on their amino acid content. Training the model on public epitopes, we discovered that the majority of public linear epitopes span about 15 amino acids and that side-chain information appears to be the most influential factor for distinguishing peptides with and without epitopes. Our results contrast prior work that could not reliably distinguish epitopes based on amino acid sequences alone (Akbar et al. 2021). In the future, it will be important to investigate whether using outputs from more complex algorithms like Alphafold2 (Jumper et al. 2021) can improve predictions. The Dolphyn pipeline is prepared to accept new models.

Using phage display libraries designed with Pepsyn and Dolphyn, we studied the immune response of healthy individuals to gut phage. We found that the majority of individuals showed reactivity to *E*.*coli*-infecting *Myoviridae*. Both libraries captured this relationship. It remains to be determined whether these antibodies are functional (e.g. enhancing or neutralizing), and what sorts of health-related phenotypes, if any, associate with these immune responses.

In short, Dolphyn libraries require much fewer peptides to reveal key patterns in antibody reactivity, giving it an advantage over libraries that contain regularly spaced overlapping peptides. The modularity of Dolphyn, such as the interchangeability of the epitope prediction module, further highlights its potential for future applications in peptide library design for the immunological study of large proteomes, such as the entire gut, skin, or lung microbiome.

## Methods

### Peptide libraries and cohorts

#### Public Epitope Library

(for the study of the nature of epitopes and training the random forest classifier):

This T7 bacteriophage display library contains 357 56 amino acid long *wildtype* peptides that showed frequent antibody reactivity (public epitope peptides) in a previous VirScan study (Xu et al. 2015). Within each of these wildtype peptides, a series of shorter peptides of length 15, 20, 25, 30, 35, 40 and 45 amino acids were designed to tile across the original 56 amino acid peptide in steps of 5 amino acids (Fig 1B). In addition, each of the wildtype peptides was subject to triple alanine mutation scanning as described previously (Xu et al. 2015). This library contains 46,070 peptides.

#### GPD Phageome Pilot Library

(for evaluating the performance and demonstrating the utility of the Dolphyn algorithm):

This library contains three subsets of peptides, representing the same 112 prevalent phages in 3,354 protein cluster representatives:

1. 19,117 peptides (length = 15 amino acids) that are likely to contain an epitope based on the random forest predictions (value > 0.5). These encoding oligonucleotides are padded on the 5’ end to make them the same length as the other two peptide libraries (56 amino acids), with three stop codons and a random sequence generated with a pseudo-random generator, i.e. the Python random.choice() function.
2. 5,266 peptides designed with the Dolphyn algorithm. 15-mer epitope peptides are grouped if they are present on several proteins. A *Dolphyn peptide* is created for every three epitope 15-mers, that are available per protein group. The 15-mer having the highest-probability epitope goes first, then a GGGGS linker, then the 15-mer having the second highest probability, then a GGGGS linker, then the 15-mer having the third highest probability, then a stop codon, creating a peptide of 56 amino acid length. If two or more Dolphyn peptides are created per protein set, the second highest probability 15-mer gets the first position on the second peptide and all other epitopes are ranked and positioned accordingly.
3. 23,745 Pepsyn peptides created by tiling the protein sequence with 56 amino acid long peptides and overlapping by 28 amino acids. If the protein length is not a multiple of 28, a full 56-mer is created at the C-terminus of the protein, potentially overlapping more than 28 amino acids of the previous tile.

All three sub-libraries were reverse translated with the Python Pepsyn package’s revtrans command to obtain 168-nt long oligonucleotides. A 16-nt long prefix (AGGAATTCCGCTGCGT) and suffix (ATGGTCACAGCTGTGC) were added to each oligo for PCR amplification, making the oligonucleotides 200-nt long. The final Dolphyn library is comprised of 48,128 individual 200-mers. The oligonucleotide library was synthesized by Twist Bioscience (San Francisco, CA).

#### Enterovirus Sublibrary

(for evaluating the prediction performance of the random forest classifier):

This library contains 1,514 peptides derived from seven reference enterovirus sequences. The sequences were selected using the cd-hit tool to represent each species of Enterovirus A, B, C, D and Rhinovirus A, B, C. Based on the selected sequences, we designed two sets of peptides using Dolphyn and Pepsyn algorithms. The first half of peptides consists of 757 epitope peptides with a length of 15 amino acids that were selected using the random forest scoring method. The second half was generated by adding a stop codon to the C-terminus of the first half of epitope peptides.

To convert the designed amino acid peptides to oligonucleotide sequences of uniform length, the Pepsyn algorithm was employed. The function ‘revtrans’ reverse-translated the amino acids sequences into DNA sequences by randomly choosing codons based on the *E*.*coli* codon usage table, with a frequency threshold of 0.1. Since the designed peptides varied in length, they were padded to a length of 120nt with a linker sequence, GCAAGTCCTGCAGCTCCAGCCCCTGCAAGCCCAGCAGCTCCAGCACCAAGTGCACCTGCTGGCGGAGGAGGTTCTGGCGGGGGCGGGAGC. Prefix AGGAATTCCGCTGCGT and suffix GTCGTGACTGGGAAAC were added for cloning purposes. Pepsyn’s ‘recodesite’ command was used to eliminate all EcoRI (GAATTC) and HindIII (AAGCTT) sites in the oligonucleotide sequences, as they were used to clone the library inserts into the T7 vector. The oligonucleotide library was synthesized by GenScript Biotech (Piscataway, NJ) using their oligonucleotide library synthesis platform.

#### Enterovirus library screening cohort

(used with the Enterovirus Sublibrary for evaluating RF prediction performance):

The Diabetes Autoimmunity Study in the Young (DAISY) (clinicaltrials.gov identifier: NCT03205865) is a longitudinal study of children at high risk for development of Type 1 Diabetes (T1D) due to genetic markers or family history. The cohort comprises approximately 7% African American, 30% Hispanic, and 63% non-Hispanic white, with the remaining participants being of biracial or other ethnicity. The study follows participants from birth, collecting blood samples annually for autoantibody testing and other biological samples for future analysis. If T1D-related autoantibodies are found, the subjects are closely monitored for the onset of the disease. The DAISY cohort was recruited between 1993 and 2004, and follow-up data is available until February 2018 (Monaco et al. 2022). To conduct a PhIP-Seq screen with the Enterovirus sublibrary, a total of 55 patient samples were selected from a subset of six subjects, consisting of three T1D positive and three T1D negative individuals.

#### VRC cohort

(used with the GPD Phageome Pilot Library, and previously with VirScan (EBV results):

The Vaccine Research Center (VRC) cohort has been described previously (Venkataraman et al. 2022) and is comprised of 801 healthy community volunteers in the greater Baltimore/Washington DC area recruited for research studies. Of the 801 individuals, 535 are of European genetic ancestry, 194 of African genetic ancestry, 32 of Asian genetic ancestry and 40 belonging to other ancestral groups. The VRC cohort included 446 men, 351 women and 4 “unknown”. Their ages ranged from 18-70 years, with an average of 35.79 years. VirScan was performed on all 801 VRC subjects. A subset of 50 subjects were selected for a PhIP-Seq screen with the GPD pilot library set. Of the 50 individuals, 41 are of European genetic ancestry, 4 of African genetic ancestry, 4 of Asian genetic ancestry and 1 belonging to other ancestral groups. The group consisted of 22 men and 28 women between ages 18 and 64, with an average age of 39.

#### PEDS cohort

(used with the Public Epitope Library to create the Public Epitope DataSet (PEDS)):

425 plasma samples (Table S1) for public epitope testing were obtained from the Genital Shedding (GS) Study (Uganda and Zimbabwe; 2001-2009), which evaluated the relationship between hormonal contraceptive use, genital shedding of HIV, and HIV disease progression among women with known dates of HIV seroconversion (Morrison et al., 2011). ART was recommended for study participants with CD4 cell counts below 250 cells/mm3, consistent with local treatment guidelines at the time the GS Study was performed. Data for CD4 cell count and viral load were collected in the GS Study (Morrison et al. 2011) data on the timing of ART initiation was obtained by review of clinic records.

### Experimental Methods

#### Library construction

Each oligonucleotide pool was resuspended in ultrapure H_2_O to a concentration of 10 ng/μl. A first round of 2 cycles of PCR was performed using 1ng of library DNA and the primers GCGCAAATGGGCGGTAGGCGTGAGGAATTCCGCTGCGT (forward) and GATTAACCCTCACTAAAGGGAAAGCTTGCACAGCTGTGACCAT (reverse). The PCR product was purified and a second round of 12 cycles of PCR was performed on all recovered PCR product with the primers CGCAAATGGGCGGTAGGCGTG (forward) and ATTAACCCTCACTAAAGGGA (reverse). The amplified DNA was purified using a PCR purification column, digested with EcoRI and HindIII, gel purified and ligated with EcoRI/HindIII digested T7FNS2 vector arms. (Shrock, Shrock, and Elledge 2022) The ligated inserts were packaged with the T7Select packaging kit (Millipore Sigma, St. Louis, MO) as per manufacturer’s instructions. An adequate number of packaging reactions to ensure a 100X coverage of the library were set up, pooled and a “pre-amplification” phage stock was prepared by the plate amplification method. The pre-amplification phage library stock was titered, mixed with DMSO at a final concentration of 10% and stored at –80 °C for the long term. A “post-amplification” working stock of the library was prepared for PhIP-Seq, using the “liquid” amplification method, ensuring that a minimum plaque forming units (pfu) of at least a 100X the library size was used as input. The post-amplification library was titered and a pfu of 100,000X the size of the library was mixed with each sample for PhIP-Seq.

#### Phage Immunoprecipitation Sequencing (PhIP-Seq)

PhIP-Seq was performed as previously described. (Mohan et al. 2018) Briefly, 0.2 μl of serum sample was incubated with the phage library overnight at 4 °C. The serum-phage mixture was then incubated with a mixture of 20 μl magnetic protein A beads and 20 μl protein G beads for 4 hours to immunocapture serum IgG antibodies and antibody-bound phage. The phage-antibody complexes captured on the beads were washed to remove unbound phage. After bead washing, peptide-coding DNA inserts from the phage were PCR amplified with forward and reverse primers containing dual indexed adapters suitable for Illumina sequencing. The PCR amplicons were pooled and subjected to DNA sequencing on an Illumina NextSeq 500 instrument.

### Computational Methods

#### Preliminary analysis of PhIP-Seq data

The output from short-read sequencing of the immunoprecipitated phage libraries underwent initial processing using the edgeR pipeline as previously described (Chen et al. 2022). Sequencing reads of a phage library member were counted via exact matching of the first 50 peptide coding nucleotides and a pseudocount was added. The magnitude of reactivity to each library member relative to mock immunoprecipitations, as defined by a fold change and associated p-value, was determined using the edgeR package in R (Robinson, McCarthy, and Smyth 2010). Significant reactivity or “hits” were defined as library members with counts greater than 15, fold change greater than five, and p value less than 0.001 (referred to as hit foldchange, “hfc”, throughout this manuscript). All other analysis of PhIP-Seq data was performed subsequent to this initial processing.

#### Phage Aggregate Reactivity Score (PhARscore)

To facilitate interpretation of complex antibody reactivity profiles, we modified the ARscore algorithm to aggregate antibody reactivity to all peptides that represent each phage (Morgenlander 2023). The GPD contains many homologous proteins, resulting in peptides that represent multiple proteins and phages. Phage-association of each peptide was tracked at every clustering step during the library design. Phage aggregate reactivity scores (PhARscores) were calculated for each phage represented by ≥ 25 peptides in a sublibrary (112 phages in the Pepsyn sublibrary, 88 phages in the dolphyn sublibrary). PhARscores from each sublibrary were generated separately.

PhARscores for a given phage were calculated by comparing mean log2 foldchange of each phage-associated peptide set to distributions of mean log2 foldchange values of the same number of randomly drawn peptides from the same sublibrary. This process was then repeated whereby, in each iteration peptides from strongly reactive phage (PhARscore > 1) were removed from the pool of peptides used to generate random distributions. This process was performed a maximum of seven times or until no new phage met the reactivity threshold.

#### ML training set

The 15-mers of the PEDS serve as training data. A 15-mer from the PEDS dataset is considered to be reactive if 2 or more samples have a log(hit-foldchange) > 0. For negative examples, we choose 15-mers with high count on the (empty) bead samples with no reactivity in any sample, to avoid including sequences that may not show reactivity due to technical reasons. The dataset was constructed to have the same number of positive and negative examples (balanced dataset). The advantage of this dataset is that many similar training examples (sequences) are contained with differing labels. When splitting test- and training-set (5/95%), we ensured that sequences derived from the same wildtype were not present in both sets.

#### Random Forest Classifier

For binary classification of 15 amino acid long peptides, a Random Forest model (Python scikit-learn package version 0.24.2 *RandomForestClassifier*) was trained on 556 features (Fig 2D). Default values for model parameters were used, including the number of trees (n_estimators_ = 100) and setting the random_state = 42 for reproducibility.

A random forest model combines several features’ impact. A particular feature’s importance (impurity based) is measured and can be extracted from the model. We report the top 9 features in Figure 2C. A higher importance value indicates that the feature is more effective at distinguishing the two classes. The *RandomForestClassifier* also provides out-of-bag scores, summarizing the prediction performance of the random forest model on out-of-bag samples, which were used for Fig 2A.

#### EBV Random Forest Testing

Epstein-Barr Virus (EBV) peptides (56-mers, Pepsyn design) from the VirScan library were used to assess our model predictions. There were no EBV peptides present in the training dataset. 801 samples from the VRC cohort were used to establish a ground truth as to whether a peptide is reactive or not. A positive label is given when at least two members of the cohort showed reactivity. All sub-15-mers were evaluated with the Random Forest classifier. The mean probability of all 15-mers in the 56-mer determines the probability score for the peptide, from which the ROC curve in Figure 2B was constructed.

#### Principal Component (PCA)

The PCA (results in Figure 5E) was conducted with the base R (version 3.6.3) function prcomp (*stats* package) based on the PhARscore vector of all samples for each phage. The first two principal components were plotted and per panel colored differently for various phage meta information.

#### BLAST for additional phage annotation

To annotate phage genomes according to whether they potentially infect *E*.*coli* or other Bacteria, we used blastn version 2.13.0+ to scan the entire NCBI nt database for similarity. We considered that if a phage genome was contained in a bacterial genome in the database, the phage may have infected that bacterium and was sequenced alongside when the reference genome was created. A BLAST hit to a viral taxonomy might indicate a potential taxonomic annotation for these novel phages.

For the binary annotation in this manuscript, an assignment is used that indicates whether an *E*.*coli* genome is among the top 15 BLAST hits that have an e-value < 1E-6.

#### Metadata Analysis of the GPD

The GPD (Camarillo-Guerrero et al. 2021) contains metadata for all phages. The predicted phage taxon and predicted host was used to annotate the heatmap in Figure 4D. Furthermore, this metadata contains a list of sample identifiers, corresponding to the cohort used in the study, of whose metagenomic samples the phage genomes were derived. We counted the amount of samples annotated as “North American” to i) select the locally prevalent phages in our pilot library and ii) conduct the prevalence study in Figure 5B.

## Data Availability

**The GitHub repository** contains all data and script resources, such as the scripts for deriving the results (folder Manuscript Analyses), raw data, the Dolphyn python package and the ML models as described in the manuscript. It is available at https://github.com/kepsi/Dolphyn and on **Zenodo** (https://doi.org/10.5281/zenodo.7979557).

## Acknowledgements

We thank the members of the NIH Vaccine Research Center for healthy volunteer serum sample collection: M. Roederer, B. Graham, L. Novick, J. Casazza, J. Ledgerwood, U. Sarwar, L. Chang, C. Starr Hendel, L. Holman, S. Plummer, P. Costner, I. Gorden, B. Larkin, F. Mendoza, J. Saudners, K. Zephir, M. E. Enama, G. Yamshchikov, I. Pittman and P. Williams.

