## Supplementary Material for "Efficient encoding of large antigenic spaces by epitope prioritization with Dolphyn"

##### Figure S1 Dolphyn Algorithm in Detail

Dolphyn predicts for each 15-amino-acid sub-peptide (15-mer) of a protein sequence whether it contains an epitope using our random forest model described in the manuscript. Hence, a prediction profile is created as displayed on the right. The 15-mer with the highest probability is chosen first and all 15-mer predictions that overlap with this first 15-mer are masked out before selecting the next highest 15-mer prediction. The masking and selecting is repeated until there are no more 15-mers with prediction values greater than the cut-off (default 0.5).

For creating the stitched peptides, the highest probability 15-mers are chosen in a number that is devisable by three. For example, if a protein sequence contains 7 potential epitopes, 6 of them are used for stitching onto 2 peptides that represent this protein.

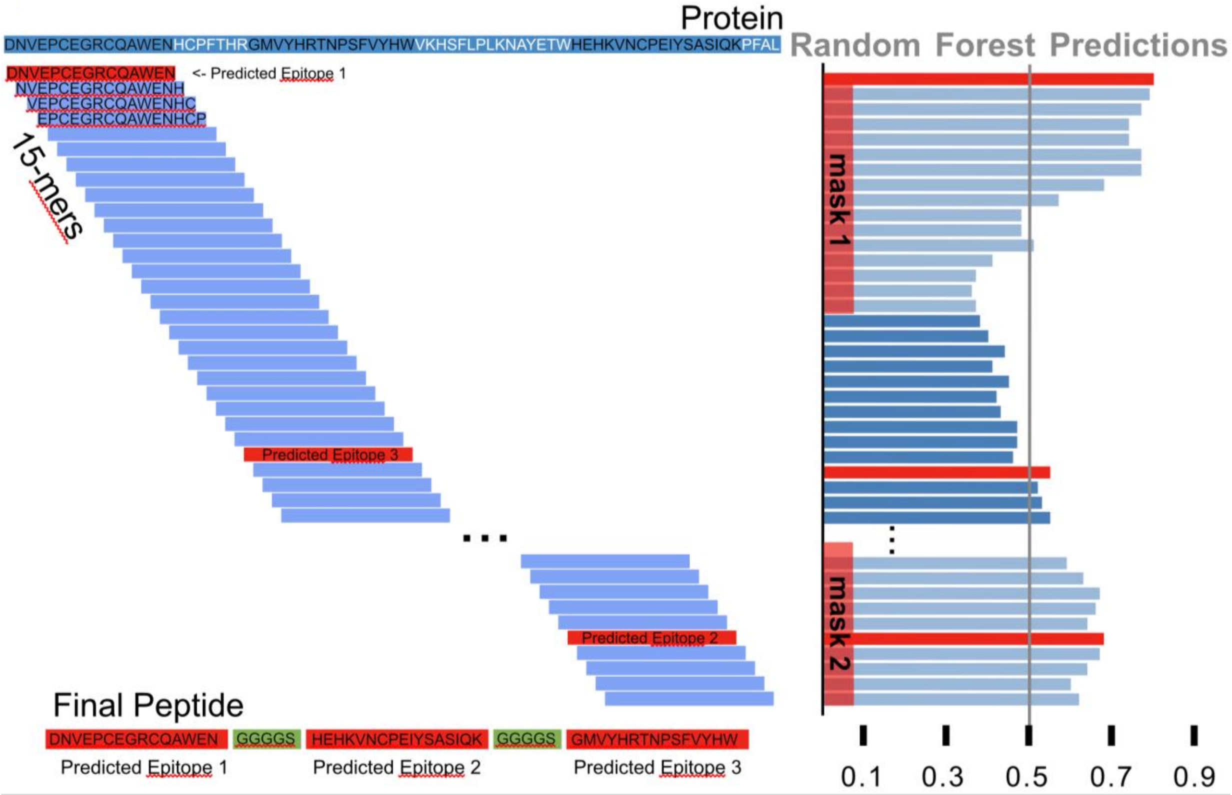

### Figure S2 Linker Sequences

In Dolphyn's "stitching" step, three 15 amino acid long peptides are combined onto one peptide and separated via a flexible and inert linker sequence GGGGS. We evaluated two different linkers, a single Glycine (G) or four Glycine and one Serine (G4S = GGGGS). We found that the G4S linker allows a higher reactivity value (hit foldchange) for the same peptides.

A validation library was created from the PEDS for evaluating the stitching approach and two different linker sequences between the epitopes. Having ground truth for these 15 amino acid long peptides, we include the 48 most reactive ones, i.e., the ones reactive in the highest number of tested samples, into this validation library. Further, we included 96 "non-reactive" peptides in the library which are those that have high bead counts in the quality control samples, but no reactivity in in the tested patients. We ensured, not to select peptides, reactive or not, from the same wildtype. Then, we stitched the 48 reactive peptides onto 16 tiles with three epitopes, each and 96 non- reactive one onto 32 tiles. Next, we combine reactive and non-reactive peptides randomly with three peptides per tile. All stitched peptides will be synthesized twice, with the two different linkers. This library totals 336 peptide tiles. Any sequence shorter than 56 amino acids was filled up with three stop codons and a random sequence of DNA nucleotides.

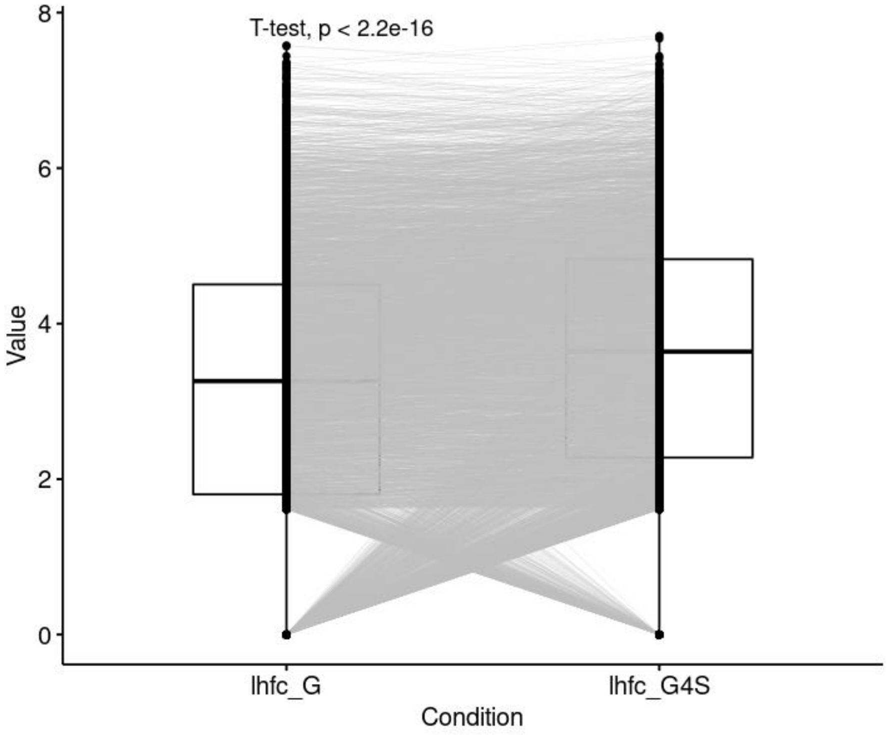


Figure S2 - Paired t-test of triplet peptides separated by G or GGGGS linker.

### Figure S3 Protein Discovery Power of Library Peptides

Figure 5D in the manuscript shows the performance metrics recall, precision, F1 and accuracy for Dolphyn-library and Pepsyn-library peptides. All peptide-sample pairs are compared to the ground-truth. The ground-truth in Figure 5D is defined from the Pepsyn library. A positive hit is defined, if any peptide in a protein is reactive in an individual. The performance metrics then are calculated for all peptides in the library (which is why Pepsyn's recall is not 1).

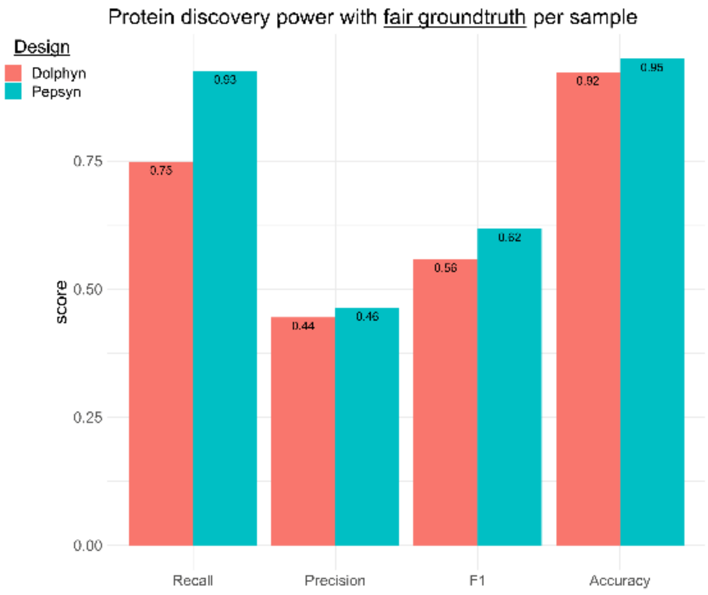


Figure S3 – The ground-truth is defined if any peptide of both libraries, Pesyn or Dolphyn, in that protein displays reactivity in an individual.

### Table S1 Cohort for Public Epitope Dataset

439 plasma samples were obtained from the Genital Shedding (GS) Study (Uganda and Zimbabwe; 2001-2009) for the creation of the public epitope dataset (PEDS). Table S3:

| **Characteristic** | **Cohort** |
|---|---|
| Country of origin | Zimbabwe |
| HIV subtype | C |
| Number of samples | 425 + 14 replica |
| Number of participants | 59 |
| Duration of HIV infection (years) | 0.04 to 8.7 |
| Mean samples per participant, (range) | 7 (3 - 9) |
| Female sex, % of participants | 100% |

### Available on GitHub: Public Epitope Visualization

The GitHub repository features a PDF which visualizes each wildtype contained in the public epitope dataset individually. Each page contains a graphic, such as the following, where each line is the alanine scan for one sample. In particular, the value on the y-axis in the difference in fold-change from the wildtype fold-change.

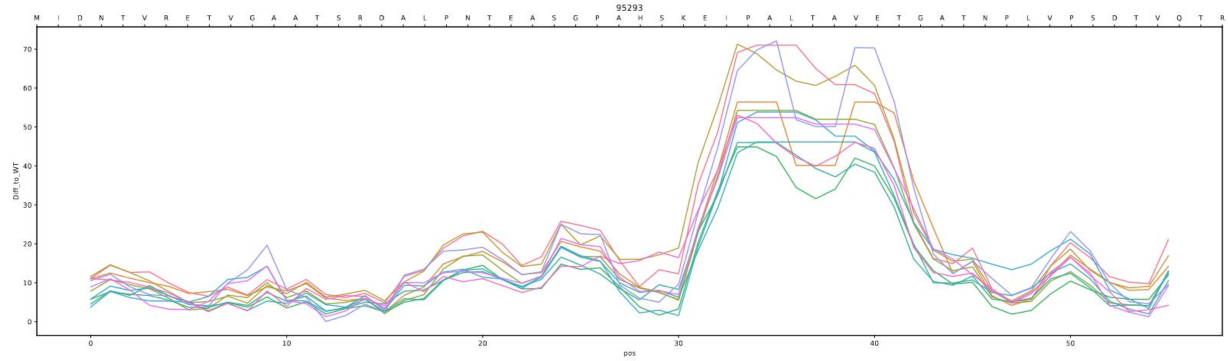


Sample page of file *AlanineScan_DiffToWT.pdf* on https://github.com/kepsi/Dolphyn .
